# Dissecting the invasion of *Galleria mellonella* by *Yersinia enterocolitica* reveals metabolic adaptations and a role of a phage lysis cassette in insect killing

**DOI:** 10.1101/2022.06.27.497489

**Authors:** Philipp-Albert Sänger, Stefanie Wagner, Elisabeth Liebler-Tenorio, Thilo M. Fuchs

**Affiliations:** Friedrich-Loeffler-Institut, Institut für Molekulare Pathogenese, Naumburger Str. 96a, 07743 Jena, Germany

**Keywords:** *Yersinia enterocolitica*, insecticidal toxin, TcaA, lysis cassette, *Galleria mellonella*

## Abstract

The human pathogen *Yersinia enterocolitica* strain W22703 is characterized by its toxicity towards invertebrates that requires the insecticidal toxin complex (Tc) proteins encoded by the pathogenicity island Tc-PAI_*Ye*_. Molecular and pathophysiological details of insect larvae infection and killing by this pathogen, however, have not been dissected. Here, we applied oral infection of *Galleria mellonella* (Greater wax moth) larvae to study the colonisation, proliferation, tissue invasion, and killing activity of W22703. We demonstrated that this strain is strongly toxic towards the larvae, in which they proliferate by more than three orders of magnitude within six days post infection. Deletion mutants of genes *tcaA* and *tccC* were atoxic for the insect. W22703 ΔtccC, in contrast to W22703 Δ*tcaA,* initially proliferated before being eliminated from the host, thus confirming TcaA and TccC as membrane-binding Tc subunit and ADP-ribosylating enzyme, respectively. Time course experiments revealed a Tc-dependent infection process starting with midgut colonisation that is followed by invasion of the hemolymph where the pathogen elicits morphological changes of hemocytes and strongly proliferates. The *in vivo* transcriptome of strain W22703 shows that the pathogen undergoes a drastic reprogramming of central cell functions and gains access to numerous carbohydrate and amino acid resources within the insect. Strikingly, a mutant lacking a phage-related holin/endolysin (HE) cassette, which is located within Tc-PAI_*Ye*_, resembled the phenotypes of W22703 Δ*tcaA,* suggesting that this dual lysis cassette is an example for a phage-related function that has been adapted for the release of a bacterial toxin.

## Introduction

Few bacteria are known to successfully colonize and infect invertebrates and to eventually profit from their bioconversion. Key factors for insect infection are the insecticidal toxin complex (Tc) proteins, which were first purified from *Photorhabdus luminescens* (1). Their oral insecticidal activity is comparable to that of the *Bacillus thuringiensis-* (Bt-) toxin. Homologues of the Tc proteins have been described in insect-associated bacteria such as *Serratia entomophila* and *Xenorhabdus nematophilus.* 3-D structural analysis of the tripartite Tc suggests a 4:1:1 stoichiometry of the A, B and C subunits, with the A subunit forming a cage-like pentamer that associates with a tightly bound 1: 1 sub-complex of B and C (2, 3). The TcaA subunits are assumed to bind to the membranes of insect midgut cells and harbour a neuraminidase-like region that possibly confers host-specificity (4). The B and C proteins form a large hollow structure encapsulating the toxic and highly variable carboxyl-terminus of TccC that has recently been demonstrated to ADP-ribosylate actin and Rho-GTPases (5–7). The attachment of the Tc to the host cell membrane is probably followed by receptor-mediated endocytosis (2, 8). A pH decrease then triggers the injection of a translocation channel formed by the pentameric TcaA subunits into the endosomal vacuole, followed by the subsequent release of the BC subcomplex into the cytosol of the target cell (4).

Insecticidal Tc proteins are also present in the three human pathogenic and in some environmental *Yersinia* species (9). The species *Y. enterocolitica* is characterized by a unique lifecycle, as some of its representatives are able to switch between two distinct pathogenicity phases that manifest in invertebrates or mammals (10). Strain W22703 (biotype 2, serotype O:9) carries the highly conserved chromosomal pathogenicity island Tc-PAI_*Ye*_ that encodes a regulator and the type A (*tcaA*, *tcaB*), type B (*tcaC*) and type C (*tccC1*) toxin complex subunits. TcaA of *Y. enterocolitica* was demonstrated to be essential for toxic activity towards larvae of the tobacco hornworm *Manduca sexta* and the nematode *C. elegans* upon oral uptake of cell lysates or living cells (11, 12). Its Tc proteins are produced at environmental temperatures, but silenced at 37°C. The thermodependent activation of insecticidal activity is mainly the result of an antagonism between the regulators TcaR2 and YmoA. The thermolabile TcaR2 is essential and sufficient to activate *tc* gene transcription at low temperatures (13), whereas the *Yersinia* modulator of virulence, YmoA, a Hha-like protein that interacts with the DNA-binding protein H-NS, represses *tc* gene transcription at 37°C (14).

Remarkably, two highly conserved phage-related genes, which are not clustered with other phage determinants, are present in all insecticidal pathogenicity islands identified so far in *Yersinia* strains (9). These genes termed *holY* and *elyY* are located between *tcaC* and *tccC* (**Figure S1**), and their products were recently shown to act as a holin and an endolysin, respectively. ElyY revealed an endopeptidase with high substrate specificity that cleaves yersinial murein. We also demonstrated that upon overexpression, the dual lysis cassette is able to lyse cells of *E. coli* and of several, but not all, *Yersinia* species including *Y. enterocolitica* and *Y. pseudotuberculosis* (15). Remarkably, HolY and ElyY lyse *Y. enterocolitica* at body temperature, but not at 15°C. When the gene encoding the Lon protease was deleted, lysis of the mutant was observed at 15°C, indicating that this enzyme controls the temperature-dependent activity of the holin/endolysin cassette (16). The biological role of this lysis cassette and its potential contribution to the insecticidal activity of *Y. enterocolitica* has not been elucidated.

Invertebrates are often used as alternative to mammalian models of infection to study bacterial or fungal pathogenicity and to evaluate therapeutic interventions (17). Nematodes, which are easily maintained in the laboratory without raising ethical issues, have successfully been used to identify virulence-related genes in a broad set of bacterial pathogens (12, 18). On the other hand, insect models such as *Galleria mellonella,* the Greater wax moth, are considered to provide further insights into pathogen-host-interactions due to their more elaborated innate immune system (19, 20). They also allow subcutaneous injection as well as oral application of bacteria and fungi, *in vivo* imaging of bacterial cells, monitoring of intracellular gene expression, detection of immune responses, and the investigation of antimicrobial drugs (21–24). Subcutaneous infection of *G. mellonella* larvae demonstrated the insecticidal activity of several *Yersinia* species and identified the enterotoxin YacT of *Y. frederiksenii* (9, 25). Moreover, many *Yersinia* genes, including those contributing to virulence, are upregulated at lower temperature (26–28), corroborating the hypothesis that invertebrates are a natural host of pathogens and might therefore have fostered their evolution (29).

Here, we established and applied oral force-feeding of *G. mellonella* larvae with *Y. enterocolitica* to study the gastrointestinal and systemic infection. Survival assays, histopathology, immunofluorescence, determination of cell numbers in time courses, transcriptomics, and infections with mutants were performed to dissect the distinct stages of *G. mellonella* infection by the bacterial pathogen as well as the roles of *tc* gene products.

## Results

### Oral infection of *G. mellonella*

We first optimized the use of *G. mellonella* larvae (**Fig. 1A**) for oral infection with *Y. enterocolitica.* Bacteria injected into the mouth of larvae are expected to pass along the digestive tract (**Fig. 1B**) that they colonize to prevent excretion. Histological analysis of larvae of approximately two cm length revealed distinct sections of the digestive tract (**Fig. 1C-H**): mouth, esophagus, crop, intestine, rectum and anus. The foregut is divided from the midgut by the stomadeal valve. The midgut, the largest part of the digestive tract, is lined by glandular epithelium separated from the ingesta by the peritrophic matrix and peritrophic membrane. The glandular epithelium ends in a short transition zone. In this region the Malpighian tubes lead into the intestine, and their secretion pushes the ingesta countercurrent, in cranial direction along the glandular epithelium. The midgut ends at the proctodeal valve. The following hindgut is lined by cuticular epithelium.

**FIG. 1.**
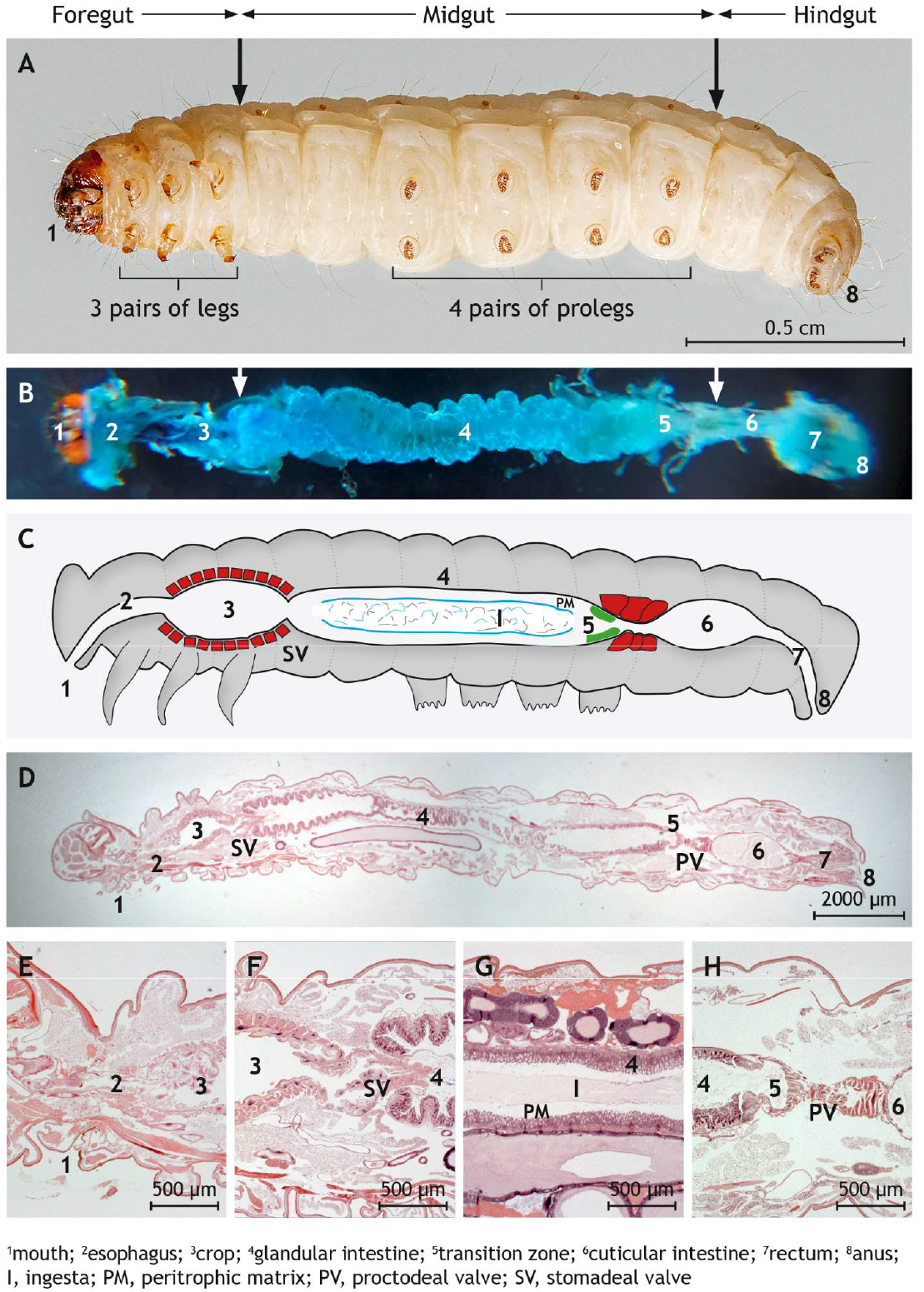
Anatomy and histology of *G. mellonella* larvae with emphasis on the digestive tract. (A) Intact *G. mellonella* larva. The beginnings and the ends of the foregut, the midgut, and the hindgut are indicated by arrows. (B) Dissected digestive tract after instillation with methylene blue. (C) Schematic drawing of the gastrointestine. The stomadeal valve (SV) is the transition from foregut to midgut, and the proctodeal valve (PV) that from midgut to hindgut. The crop and both valves are surrounded by a thick layer of musculature (red). The specific epithelium of the PV transition zone is labeled in green. The ingesta (I) in the midgut is covered by the peritrophic membrane (PM). The ingesta is separated from the mucosa by the ectoperitrophic space. (D) Longitudinal and sagittal section along the middle through *G. mellonella*. (E) Magnification of mouth, esophagus, and crop lined by the cuticular epithelium and surrounded by muscle cells. (F) Magnification of the SV between crop and glandular intestine. (G) Magnification of the glandular intestine lined by mucosa epithelium. The ingesta is surrounded by the peritrophic membrane (PM) and separated from mucosa by the ectoperitrophic space. (H) Magnification of the proctodeal valve (PV) between midgut and the cuticular intestine lined by cuticular epithelium. The lining of the digestive tract close to the PV is formed by a layer of highly vacuolated epithelial cells distinct from the glandular epithelium of the midgut. (D)-(H) are paraffin sections stained by hematoxylin and eosin. Sections are indicated by numbers. Photos of representative preparations are shown; the scales are indicated.

Force-feeding of bacterial cultures was carefully performed by injection with a Hamilton syringe. In case of accidental tissue perforation, resulting in direct injection of bacteria into the hemacoel and thus early death of larvae, the larvae were excluded from the experiment. Due to the larva weight of 150-200 mg, a maximum of 5 μl culture or medium were applied. Preliminary experiments to establish the optimal infection dose were performed with a range of 10^2^ to 10^8^ colony forming units (CFU) of *Y. enterocolitica* and showed dose-dependent phenotypes. When 10^7^-10^8^ CFU were injected, all larvae died between four and 14 h p.i., and the hemolymph of these cadavers was found full of *Y. enterocolitica* cells. No lethality and no melanisation, however, was observed after applying a lower dose of 10^2^ to 10^4^ cells at least until nine days p.i., and no *Y. enterocolitica* cells could be detected in the hemolymph of the larvae. Finally, 10^5^-10^6^ CFU revealed as optimal dose to perform infections of *G. mellonella* larvae with *Y. enterocolitica*, a value that corresponds well with those used in subcutaneous applications (22, 30).

### TcaA and the lysis cassette are essential for the toxic activity of W22703 against G. *mellonella* larvae

Survival assays were performed with larvae of *G. mellonella* to further investigate the function of Tc-PAI_*Ye*_ determinants in the interaction of *Y. enterocolitica* with insects. Recently, we demonstrated that subcutaneous infection of *G. mellonella* larvae with W22703 results in a killing rate similar to that of W22703 Δ*tcaA*, suggesting that the Tc does not play a role during systemic infection (9). Here, we orally infected larvae with 5.7 × 10^5^ CFU, 7.8 × 10^5^ CFU, 5.6 × 10^5^ CFU, and 6.2 × 10^5^ CFU, respectively, of W22703 and its mutants W22703 Δ*tcaA*, W22703 ΔHE, and W22703 ΔtccC, and monitored the animals for nine days. Larvae infected with *Y. enterocolitica* strain W22703 exhibited a significantly reduced survival rate with a time to death of 50% (TD_50_) = 3.67 ± 1.12 days. In the infection experiments performed with the three mutants lacking *tcaA,* HE, and *tccC*, all larvae survived, a finding that corresponds to a 100% survival of larvae exposed to LB medium (**Fig. 2**). To genetically validate that *tcaA*, HE, and *tccC* are essential for the toxicity of W22703 towards *G. mellonella*, we orally infected larvae with W22703 ΔtcaA/pACYC-tcaA (9.0 × 10^5^ CFU), 4.0 × 10^5^ CFU (W22703 ΔHE/pACYC-HE), and W22703 ΔtccC/pBAD-tccC (4.0 × 10^5^ CFU). Due to the slight leakiness of the pBAD-promoter, *tccC* transcription was not induced by arabinose.TD_50_ of 2.91 ± 1.46 days, 1.83 ± 0.51 days, and 3.90 ± 0.41 days, respectively, were determined. Thus, the mutants harbouring recombinant plasmids that complement the deletion did not significantly differ in their insecticidal activity from that of the parental strain W22703 (TD_50_ = 3.67 ± 1.12 days), demonstrating that the *in trans* complementation of Δ*tcaA*, ΔHE, and Δ*tccC* restores the insecticidal phenotype of W2703. Taken together, these life span assays indicate that the two toxin subunits TcaA and TccC as well as the HE lysis cassette are strictly required for the toxicity of *Y. enterocolitica* strain W22703 towards the insect larvae.

**FIG. 2.**
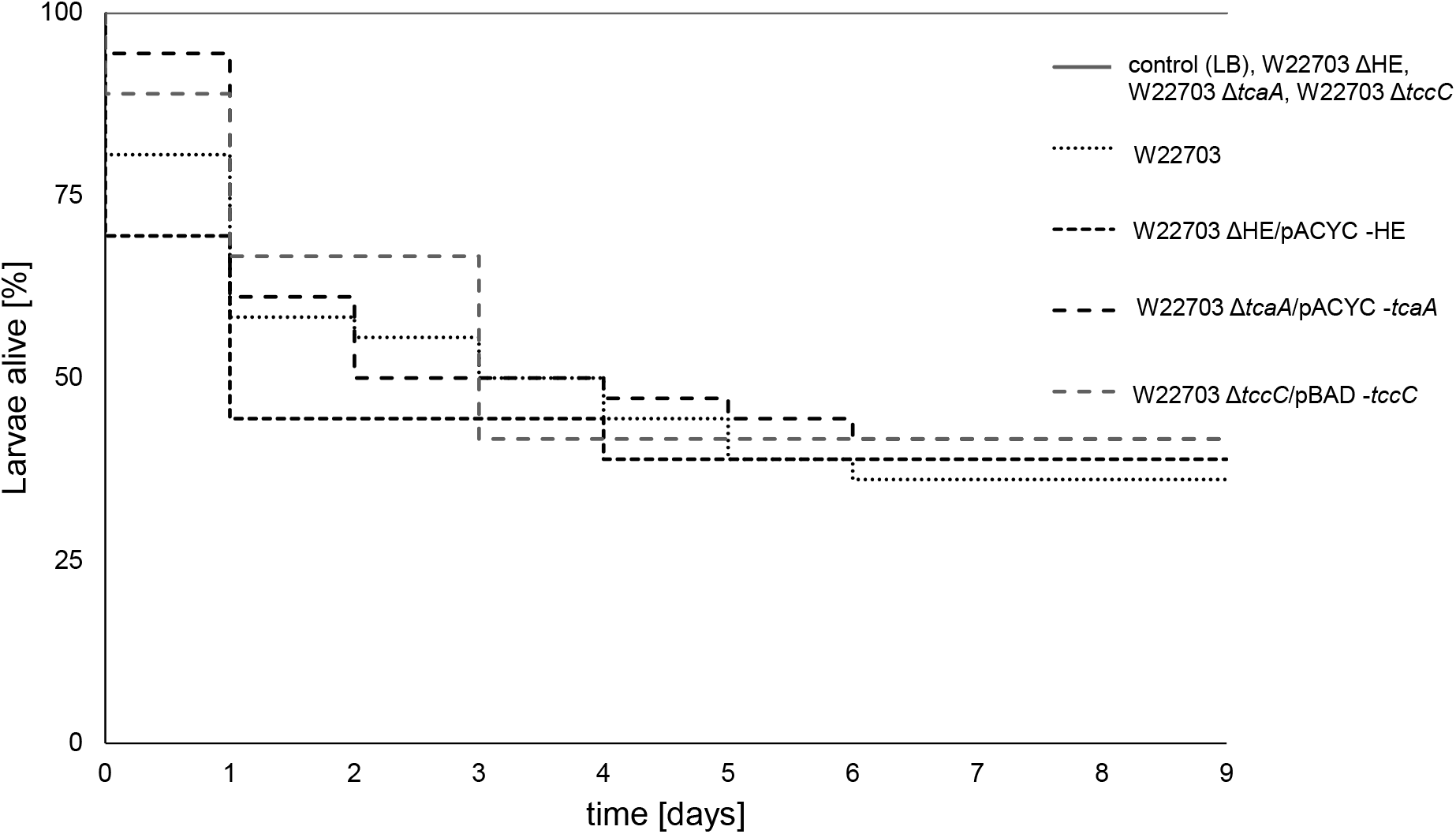
Role of TcaA, HE and TccC for insecticidal activity of W22703 towards *G. mellonella*. Larvae were orally infected with W22703, its mutants lacking *tcaA*, HE, and tccC, and with mutants carrying complementing plasmids pACYC-HE, pACYC-*tcaA*, and pBAD-*tccC*. Life span assays were performed for nine days, and the viability of the larvae was monitored each day. LB medium served as control. The raw data were plotted by the Kaplan-Meier method. The survival rate of the larvae was determined daily until day nine p.i. The Kaplan-Meier-plot is based on triplicates with 36 larvae in total per strain. The curves were compared using the log-rank test, which generates a *p* value testing the null hypothesis that the survival curves are identical. Data were fit to exponential distribution. *p* values of 0.05 or less were considered significantly different from the null hypothesis (*p* value pACYC-HE 0.0194; *p* value pACYC-tcaA 0.0369; *p* value pACYC-HE 0.0251).

In addition, we monitored the behaviour and the morphology of the larvae each day until six days post infection (p.i.), and again at day nine p.i. (**Fig. 3**). Immediately after oral application, the control group infected with LB medium did not differ from the larvae infected with the two deletion mutants W22703 ΔHE and W22703 Δ*tcaA* with respect to motility and colour. In contrast, the application of W22703 and strains W22703 ΔHE/pACYC-HE and W22703 Δ*tcaA* /pACYC-*tcaA* resulted in a higher activity of the larvae and a slight colouring of some of the larvae from one h p.i. on (data not shown). After 24 h, a strong melanization in the groups infected with W22703 and with the mutants harbouring complementing plasmids was observed. Similar to the untreated control groups, cocoons surrounded larvae that had been infected with the deletion mutants, indicating that healthy individuals only are able to produce this protective housing. At days two to three, cocoon formation continued, whereas an increasing number of larvae infected with W22703, W22703 ΔHE/pACYC-HE and W22703 Δ*tcaA /pACYC-tcaA* died. At days five to nine, morphological signs of infection did not further enhance. Nine days after infection, the first pupations events were observed.

**FIG. 3.**
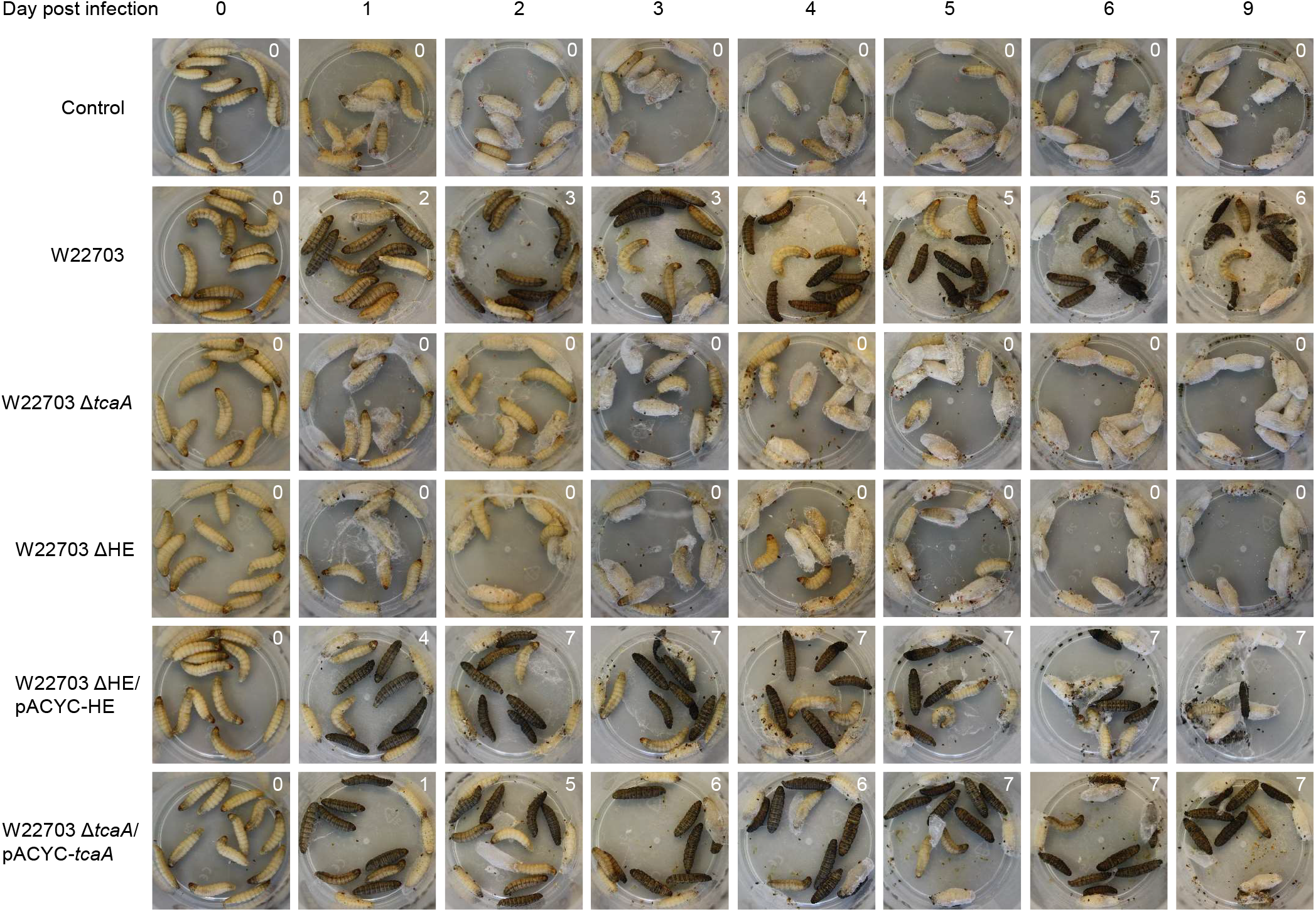
External morphology of larvae following oral infection with W22703 and its mutants. Black animals were dead, anthracite ones still alive. The numbers in the upper right angle of each photo indicates dead animals at the respective time-point. The photos were taken from the experiment shown in Fig. 2.

### Orally injected W22703 mutants lacking TcaA, HE, or TccC are eliminated by G. *mellonella*

Proliferation within the insect host would indicate a successful infection by *Y. enterocolitica*. To determine the bacterial load over the time, we infected *G. mellonella* larvae with the same strains used in Fig. 2 and applied similar infection doses. One, three and six days p.i., the homogenate of six animals per time-point were plated on selective LB agar plates, and the CFU were enumerated. Strikingly, when we injected 9.0 ± 0.2 × 10^5^ CFU of mutant W22703 Δ*tcaA*, the total numbers of surviving bacteria rapidly decreased to 1.0 × 10^3^ after one day and to 11 CFU after three days, and the strain was completely eliminated six days p.i. (**Fig. 4**). In contrast, the mutants W22703 ΔHE and W22703 Δ*tccC* initially proliferated from an application dose of 4.0 × 10^5^ CFU and 4.0 × 10^5^ CFU, respectively, to 2.2 × 10^6^ CFU and 2.8 × 10^6^ CFU, but could not be detected from day three on. This finding strongly suggests that TcaA is involved in adherence to epithelial cells and thus in midgut colonization. When larvae were infected with 9.0 × 10^5^ CFU of the parental strain W22703 and 4.0 × 10^5^ CFU of the two Δ*tcaA* and ΔHE deletion mutants carrying complementing plasmids, the bacterial burden increased to approximately 2 × 10^8^ CFU after one day and to approximately 1 × 10^9^ CFU three and six days p.i. Strain W22703 ΔtccC/pBAD-tccC of which 1.4 × 10^6^ CFU were applied did not exceed 4.6 × 10^8^ CFU, indicating a slightly weaker growth in comparison with that of the other three strains.

**FIG. 4.**
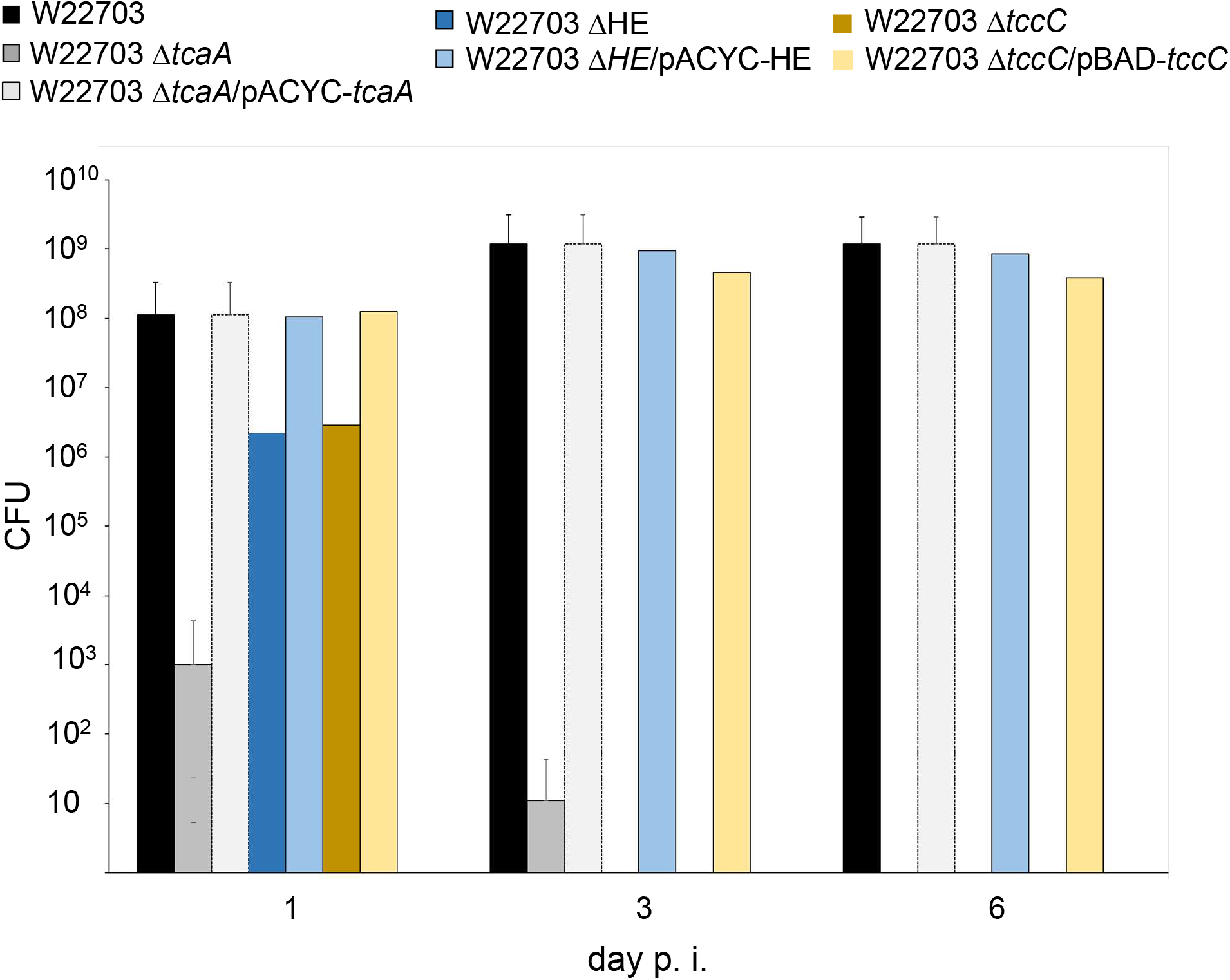
Proliferation numbers of *Y. enterocolitica* W22703 strains in *G. mellonella* larvae. Larvae were orally infected with W22703, W22703 Δ*tcaA*, W22703 Δ*tcaA*/pACAC-*tcaA*, W22703 ΔHE, W22703 ΔHE/pACYC-HE, W22703 Δ*tccC* and W22703 Δ*tccC*/pBAD-*tccC*. Six animals per time-point were used, and the experiments were performed as triplicates, resulting in a total of 18 larvae per time-point per strain. Larvae fed with LB were used as a negative control. Standard deviations are indicated as error bars.

These data show that only strains able to produce TcaA, the lysis cassette, and TccC possess the capacity to successfully colonize and to proliferate within the larvae.

### Time course of infection

The strongest proliferation of W22703 was observed between one and three days p.i. (**Fig. 4**). We hypothesized that *Y. enterocolitica* starts its infection in the midgut and then invades the tissues of *G. mellonella* larvae to proliferate in the hemolymph. To dissect the infection process preceding this multiplication in more detail, we performed a time course experiment using larvae infected with 6.3 × 10^5^ *Y. enterocolitica* W22703 cells. Longitudinal sections through the middle of the larvae were prepared 4 h, 6 h, 12 h, 18 h, and 24 h p.i. and stained with FITC-conjugated *Yersinia*-antibodies. The histological analysis revealed that *Y. enterocolitica* is present in the midgut with increasing cell numbers until 12 h p.i. At this time-point, W22703 is mainly detected close to glandular epithelial cells of the midgut, pointing to a tropism of *Y. enterocolitica* for endodermal tissue **(FIG. 5A)**. Eighteen hours p.i., the gut appears to be empty from *Yersinia*, whereas a high number of cells is now detected in the hemolymph where W22703 proliferates with the next six hours to a high cell density. The larval tissues were completely overgrown with *Y. enterocolitica* 48 h p.i. when most animals were dead, indicating that the bacteria started bioconversion of the cadaver **(FIG. 5B)**. To verify the finding that *Y. enterocolitica* W22703 cells are mainly found in the circulating fluid, we isolated 10 μl hemolymph from larvae infected with 1.6 × 10^5^ bacteria. 24 h p.i., the animals showed clear signs of melanisation. The brownish hemolymph contained 2.3 × 10^7^ to 4.7 × 10^7^ CFU as detected on selective agar.

**FIG. 5.**
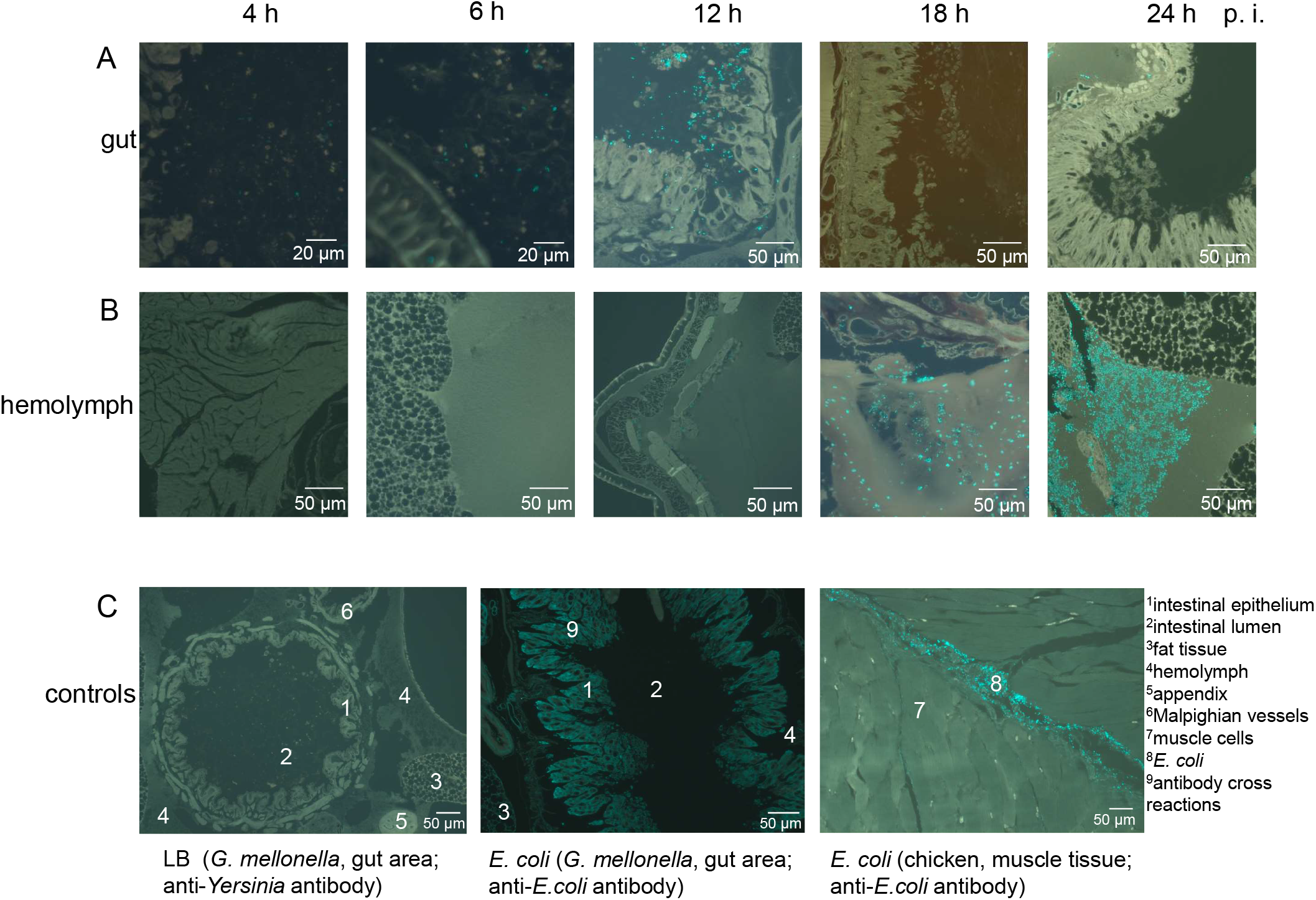
Time course of *Y. enterocolitica* W22703 proliferation in *G. mellonella*. The tissue sections monitored by fluorescence microscopy show antibody-stained *Y. enterocolitica* cells in the (A) gut or (B) hemolymph of *G. mellonella* 4 h, 6 h, 12 h, 18 h, and 24 h after infection. (C) The controls show the gut area of *G. mellonella* that were fed with LB (left) or infected with *E. coli* (middle) 24 h ago. The tissue sections were stained with a *Yersinia*-specific or an *E. coli*-specific antibody. Functionality of the anti-E. *coli* antibody was demonstrated by the application of *E. coli* into muscle tissue of chicken (right). Cyan-colored areas in the gut area of *G. mellonella* are unspecific bonds of the anti-E. *coli* antibody. Representative preparations are shown; the scale is indicated.

No fluorescence signal was obtained when we fed LB medium to larvae as a control (**FIG. 5C**). Next, *E. coli* cultures were applied analogously to the experiments with *Y. enterocolitica*, but no *E. coli* cells were detected by an anti-E. *coli* antibody in the gut or in the hemolymph of *G. mellonella* 24 h after infection. To test the specificity of the antibody, *E. coli* DH5α cells were injected into muscle cells of a chicken leg and were shown to be FITC-labeled 24 h p.i. The control experiments indicate that in contrast to *Y. enterocolitica, E. coli* is not able to survive in *G. mellonella*.

Taken together, we demonstrated that *Y. enterocolitica* systemically infects *G. mellonella* larvae *via* colonization and penetration of the midgut epithelium to finally reach the hemolymph that allows massive proliferation.

### Strains lacking insecticidal genes do not enter the hemolymph

To determine whether or not the factors encoded on the insecticidal Tc-PAI_*Ye*_ play a role during the infection process delineated above, *G. mellonella* larvae were orally infected with *Y. enterocolitica* W22703 (6.3 × 10^5^ CFU), W22703 Δ*tcaA* (8.3 × 10^5^ CFU), W22703 Δ*tccC* (6.4 × 10^5^ CFU), W22703 ΔHE (7 × 10^5^ CFU), and W22703 Δ*tcaR2* (7.8 × 10^5^ CFU). In contrast to the parental strain W22703 that proliferated to high cell numbers in the hemolymph, no mutant cells were detected by FITC-staining of tissue sections made 24 h p.i. (**FIG. 6A**). These data confirm that the insecticidal Tc as well as the *tc* gene activator TcaR2 and the HE lysis cassette are required for full virulence of *Y. enterocolitica* W22703 towards *G. mellonella*. In particular, we hypothesize that the Tc-PAI_*Ye*_ is responsible for midgut colonization and entering of the hemolymph.

**FIG. 6.**
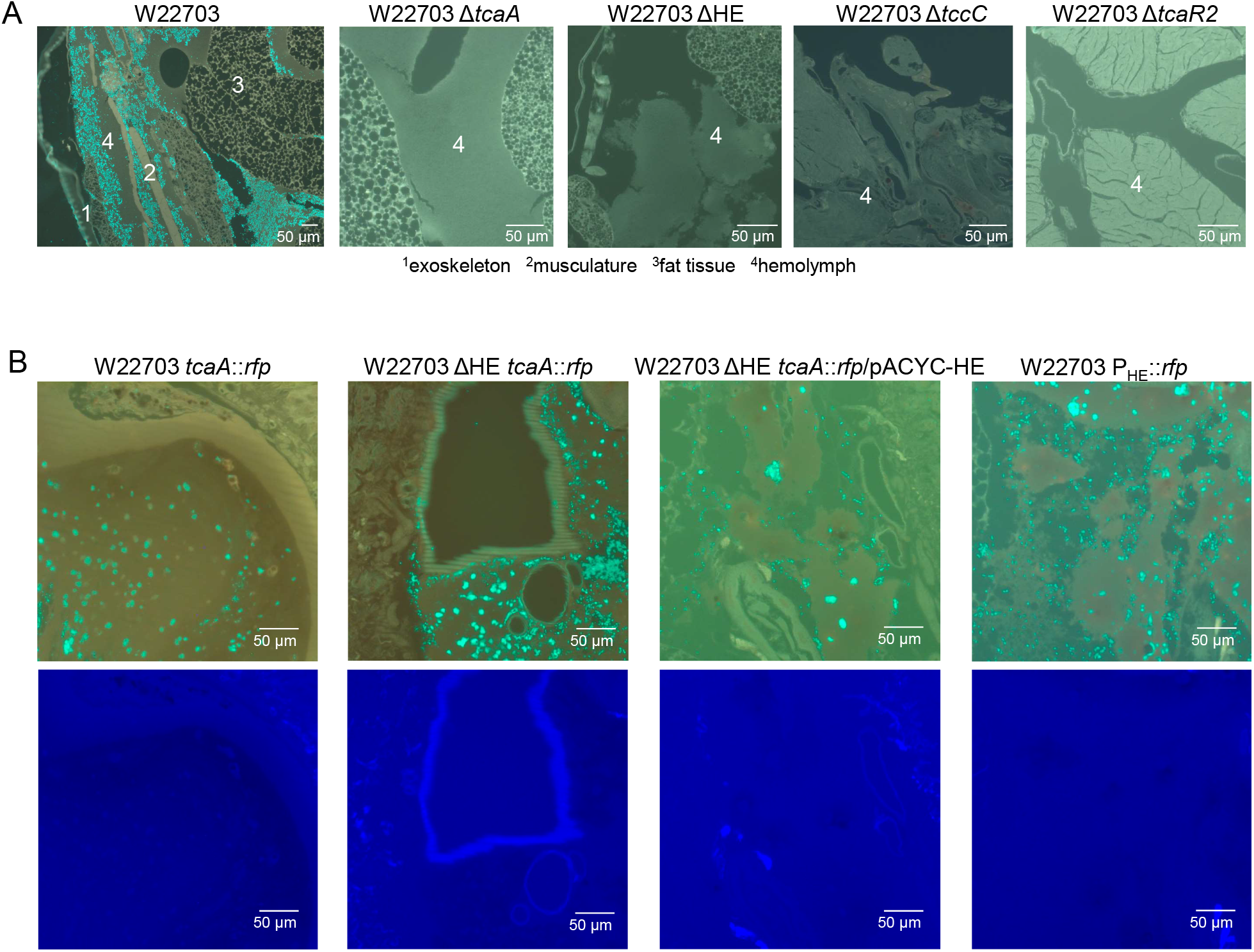
Detection of W22703 and its derivatives in the hemolymph of *G. mellonella*. (A) *G. mellonella* tissue sections 24 h p.i. with W22703 and its mutants. Staining was performed with *anti-Yersinia* antibody. (B) Tissue sections of larvae 24 h p.i. with W22703 *tcaA::rfp*, W22703 ΔHE *tcaA::rfp*, W22703 P_HE_::*rfp*, and W22703 ΔHE *tcaA::rfp*/pACYC HE. Staining was done using a *Yersinia*-specific antibody (top row) or an anti-RFP antibody (bottom row). Photos show *Yersinia* distribution of *G. mellonella* 24 h p.i. in the hemolymph area. The anti-RFP antibody was used to detect TcaA distribution in the same tissue sections. TcaA could only be detected in W22703 *tcaA*::*rfp* samples (bottom left). Preparations of 10 infected animals per strain were carried out. Photos of representative preparations are shown; the scales are indicated.

### Detection of TcaA::RFP *in vivo* depends on a functional lysis cassette

To further investigate the role of the HE cassette for the the Tc release, we infected *G. mellonella* larvae with the reporter strain W22703 *tcaA::rfp.* Following tissue section and immunostaining with anti-Yersinia antibody and anti-RFP antibody, we detected TcaA in *G. mellonella* in W22703 *tcaA::rfp* samples 24 h p.i. in hemolymph-rich areas exhibiting *Y. enterocolitica* cell cluster (**FIG. 6B**). In the absence of the lysis cassette, however, TcaA::Rfp was not detected despite the presence of W22703 ΔHE *tcaA::rfp* cells. We also failed to stain TcaA::RFP using strain W22703 ΔHE *tcaA*:: *rfp*/pACYC-HE that carries a plasmid complementing the HE deletion. This might be explained by the fact that the transcription of the holin and endolysin genes is not appropriately regulated by the pACYC construct under these conditions. To test whether or not the promoter of the lysis cassette is active *in vivo*, we infected *G. mellonella* larvae with strain W22703 P_HE_::*rfp*. Although *Y. enterocolitica* cells densely proliferated within the hemolymph (**FIG. 6B**), no staining signal that would point to the presence of TcaA was obtained, possibly due to no or weak P_HE_ activity. These data suggest that the HE cassette is responsible for the extracellular activity of the insecticidal Tc.

### Morphological effects of the TC on hemocytes

To investigate a possible activity of the insecticidal gene products on hemolymph cells, we monitored their morphology 24 p.i. For this purpose, hemolymph preparations of *G. mellonella* larvae were fixed with methanol and then stained by Giemsa solution. The hemocytes derived from W22703-treated larvae began to form aggregates in comparison with LB medium as control (**Fig. 7A**). Cell agglutinations and a fading of the chromatin color of the cell nucleus occurred more frequently. As expected, bacterial cells, e.g. *Y. enterocolitica*, are visible in the hemolymph obtained from W22703-infected animals, but not in all other preparations. In contrast, hemolymph preparations of larvae one day after oral infection with W22703 mutants lacking *tcaA*, HE, *tccC*, or *tcaR2*, which encodes the activator of *tc* genes (13), showed hemocyte cell morphologies similar to those of the untreated controls (**Fig. 7B**).

**FIG. 7.**
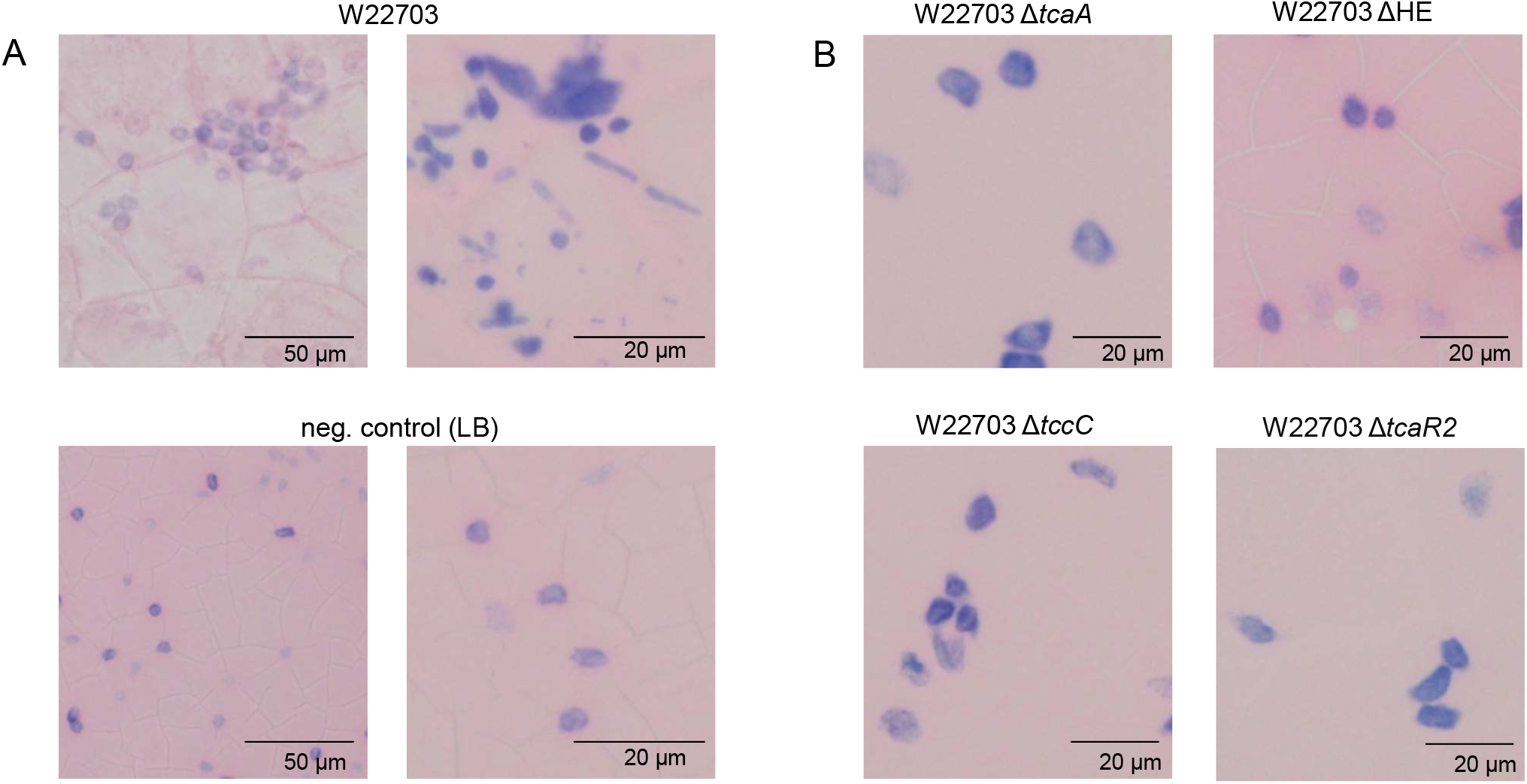
Hemocyte morphology 24 h p.i. (A) Hemolymph cell preparations from *G. mellonella* orally infected with 6.1 × 10^5^ CFU of W22703 or treated with LB medium as control. (B) Hemocytes of larvae after application of the W22703 mutants indicated, showing cell morphology similar to those of the controls. Hemocyte aggregation and deformation was visible only upon infection with W22703. Hemolymph preparations were fixed with methanol and stained by Giemsa solution. Photos of representative preparations are shown; the scale is indicated. An Olympus BX53 microscope (Olympus Europa, Hamburg, Germany) was used.

### *In vivo* transcriptome analysis

To delineate the transcriptional profile of *Y. enterocolitica* during infection of *G. mellonella*, we applied immunomagnetic separation to isolate *Y. enterocolitica* from the larvae 12 h and 24 h after infection. These two time points were chosen due to the results of the time course experiments (**FIG. 5A, B**). Growth of W22703 cells in minimal medium with glucose as carbon and energy source served as the reference condition. Following RNA sequencing and read analysis, we identified ~3,600 protein-encoding genes from a genome carrying ~4,000 genes (10), pointing to a 90% coverage of the transcriptional responses investigated here. Setting the threshold to a log2 FC ≥ |1.5|, we found 524 non-redundant transcripts to be significantly more abundant and 301 less abundant in contrast to the control (**Table S1**).

Due to their important role in the infection, colonization, and killing of *G. mellonella* demonstrated above, we first analysed the transcriptional activity of genes located on the insecticidal island Tc-PAI_*Ye*_. Strikingly, the genes encoding the activator TcaR2 and the two Tc subunits TcaA and TcaB were strongly induced (log_2_ FC = 3.9, 5.5, and 3.2, respectively) 12 h p.i., but to lesser extent 24 h p.i. (**Table S1**). The endolysin located within Tc-PAI_*Ye*_ was significantly up-regulated after 24 h, but not after 12 h, pointing to its possible role in the release of the Tc. Beside the *tc* genes, the pathogen induces 15 virulence genes including those involved in iron acquisition, whereas a set of eight is repressed, suggesting a role during infection of other host organisms including mammals.

Other main functional categories affected by a transcriptional switch upon larva infection were the metabolism and transport of amino acid and carbohydrates, resistance mechanisms, signaling, and motility (**FIG. 8**). Within larvae, *Y. enterocolitica* downregulated the biosynthesis pathways of methionine, isoleucine, leucine, histidine, tryptophan, and glutamate (**Table 1**). In contrast, determinants responsible for the transport of methionine, glycine, glutamate, glutamine, and histidine appeared at higher abundance *in vivo* in contrast to the control. This finding confirms the assumption that these amino acids are readily available in the surrounding hemolymph. The synthesis of thiamine was repressed, a finding that is in line with the reduced biosynthesis of isoleucine and leucine requiring this cofactor. Another large number of up-regulated genes belongs to the categories of carbohydrate metabolism. *Y. enterocolitica* activated transporters including phosphotransferase systems and/or enzymes involved in the uptake and/or degradation of glycerol, sorbose, mannitol, ribose, xylose, inositol, trehalose, N-acetylgalactosamine, N-acetylglucosamine, sucrose, and glucitol/sorbitol. In addition, the pathogen induced a huge set of genes encoding factors of the nucleotide and lipid metabolisms as well as the TCA-cycle, pointing to an increased metabolic activity and proliferation within insect larvae. Biofilm formation is not required for the virulence against insect as several biofilm and fimbriae producing genes were repressed, while the biofilm repressor gene was transcriptionally activated. Motility appeared to play a pivotal role in insect infection by *Y. enterocolita*, because a large set of genes involved in flagella synthesis were up-regulated. Induction and repression of genes responsible for cell membrane biosynthesis point to major rearrangements in particular of the outer membrane within the larvae. Intriguely, transcript levels of genes belonging to the *lux* regulon were reduced, while other signaling factors including the autoinducer 2 genes were transcriptionally activated in the invertebrate. No phage genes were induced, but 13 phage genes were repressed in larva-infecting *Y. enterocolitica*.

**FIG. 8.**
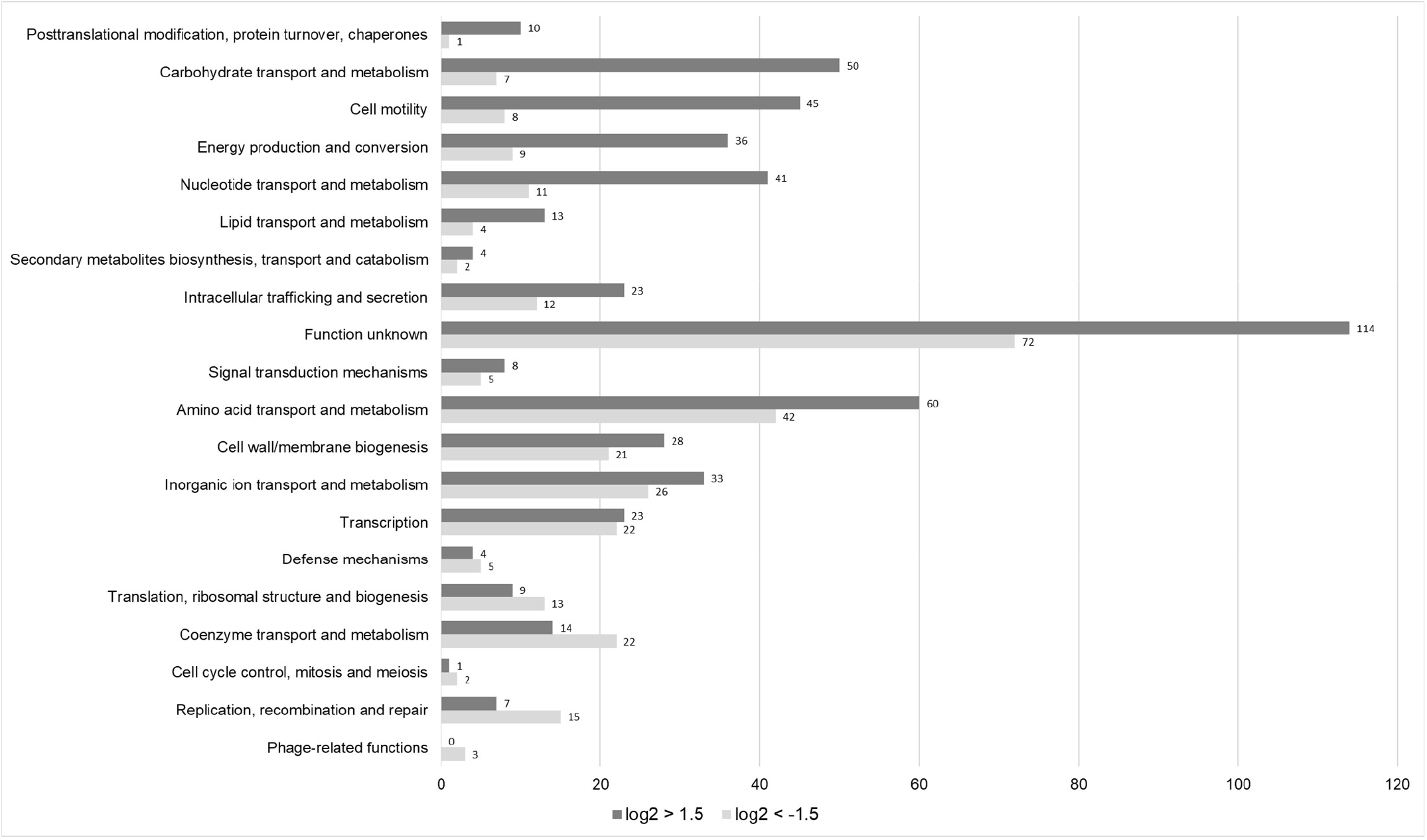
COG categories of differentially expressed W22703 genes upon *G. mellonella* infection. The numbers of down-regulated (light grey) and up-regulated (dark grey) genes in each COG category are indicated. The COG categories are ordered by descending ratio of up-to down-regulated genes. The data were taken from **Table S1**.

**Table 1:**
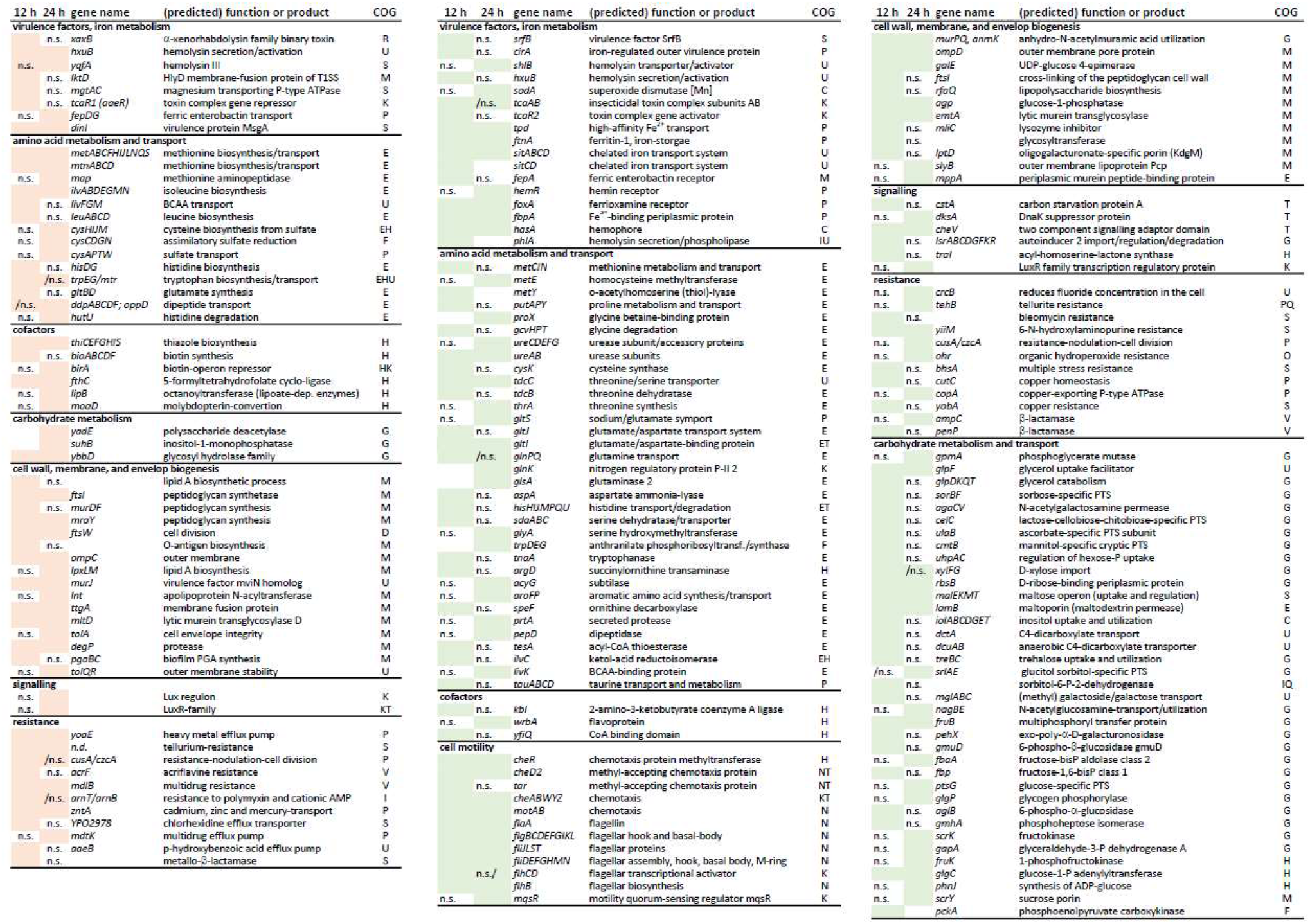
Selection of down (log_2_ FC < −1.5, red)- and up (log_2_ FC > −1.5, green)-regulated *Y. enterocolitica* genes 12 h and 24 h p.i. AMP, antimicrobial peptide; P, phosphate; PTS, phosphotransferase system; BCAA, branched-chain amino acid; dep., dependent; n.s., not significant.

Taken together, the transcriptional activity of *Y. enterocolitica* in larvae of *G. mellonella* was mainly characterized by a drastic reprogramming of the energy, amino acid and carbohydrate metabolism, by an increase of motility and signaling molecules, and by cell membrane rearrangements.

## Discussion

In order to use *G. mellonella* larvae to identify novel insecticidal factors, we subcutaneously infected its larvae with *Yersinia* species (9, 25). In the present study, we established oral application of *Y. enterocolitica* to demonstrate that *G. mellonella* larvae are a powerful tool for dissecting the steps required for a successful and lethal infection of the insect. The distinct phases of infection identified here include survival in the gut, adhesion to and penetration of the midgut epithelial cell layer, massive proliferation within the hemolymph, hemocyte deformation, and insect killing. Despite an oral infection dose between 4.0 × 10^5^ and 3.0 × 10^6^ CFU, only few *Y. enterocolitica* cells were found in the gut by cell counting and immunostaining, indicating that the majority of the bacterial cells is excreted, and that W22703 does not substantially proliferate in the gut. This phenotype is in contrast to an infection of *Manduca sexta* larvae with the entomopathogen *Photorhabdus luminescens*, during which a large fraction of *P. luminescens* cells is found to be associated with the gut (31). A few *Y. enterocolitica* W22703 cells were seen in close proximity to glandular epithelial cells of the midgut. They probably cross the epithelial barrier via M-cells and migrate into the underlying tissues. This step occurs approximately 12-18 h after ingestion, a delay reflecting the invasion process as observed also with *P. luminescens* (31). This epithelial cell contact is in line with the finding that the Tc of *Y. pseudotuberculosis* causes initial membrane ruffling of Caco-2 gut cells (32). Once they reach the hemocoel, the open circulatory system of the larvae, the pathogen exhibits a massive proliferation, pointing to excellent growth conditions and nutrient availability in this compartment. Strain W22703 seems to withstand the phagocytic or growth suppressing activities of the hemocytes, probably as a result of Tc activity as suggested by changes of the hemocyte morphology only in the presence of TcaA, TccC, and HE. To the best of our knowledge, the oral infection of *G. mellonella* by a bacterial pathogen established here is novel and of yet unprecedented resolution with respect to its readouts, although force-feeding of *G. mellonella* was already established (33, 34).

The *in vivo* transcriptome of *Y. enterocolitica* helps to better understand its physiological and biochemical adaptations to the insect. The pattern of up- and downregulated genes not only point to the relevance of distinct virulence factors, signalling, and motility for a successful infection, but also to the availability of numerous carbohydrates and amino acids in the insect body that fuels the metabolism and thus the proliferation of *Y. enterocolitica*. The carbon and energy sources are derived from either the diet or the host, such as glycerol, ribose, inositol, N-acetylgalactosamine, or trehalose, which is present in the hemolymph as well as in honey, a component of *G. mellonella* feeding. The upregulation of genes involved in the uptake and degradation of sorbose and its reduced forms glucitol/sorbitol, of mannitol, of sucrose, and of xylose point to a specific metabolic adaptation of *Y. enterocolitica* to substrates fed by insects. Sorbose and mannitol are found in plant saps, and xylose is a monomer of hemicelluloses. Sucrose is one of the most concentrated nutrient available for sap-feeding insects (35). In addition, the genes responsible for N-acetylglucosamine utilization suggests that chitin, a constituent part of the peritrophic matrices that line the inner surface of the gut in many insects (36), is not only degraded as a first step of colonization and invasion of the midgut epithelium, but metabolically utilized by the pathogen. When entering the *G. mellonella* larvae, strain W22703 in particular downregulates the genes responsible for the synthesis of methionine, branched-chain amino acids, histidine, tryptophane, cysteine, and glutamate, indicating a sufficient availability of these amino acids within the insect. This is well in line with an increased capacity to import and degrade methionine, proline, glycine, urea, cysteine, threonine, glutamate/aspartate, serine, and histidine. Histidine is one of the most abundant free amino acids in the *Hyalophora gloveri* fat body (37). It is worth to note that urease, inositol, and histidine degradation belong to those metabolic properties that are common to *Y. enterocolitica* and *P. luminescens* (28). Other examples are the up-regulated genes *dctA* and *uhp* involved in the uptake of C4-dicarboxylates and hexose-phosphate. The low temperature-dependent transcription of these and many other factors (27) corroborates the assumption that they contribute to the proliferation of W22703 in insect larvae. In addition, reprogramming the activities of *Y. enterocolitica* within insects includes the increase of lipid import and degradation, and of energy production and conversion. The latter category reflects the massive proliferation in the hemolymph. The *in vivo* transcriptional pattern also revealed the up- and downregulation of virulence factors mainly involved in hemolysis and iron scavenging. These data suggest that *Y. enterocolitica* harbours a set of determinants that are specifically directed against insects. Autoinducer-2 import was also found to be upregulated during insect infection. Given that more than 300 AI-2 regulated genes involved in regulation, metabolic activity, stress response and pathogenicity are known in *P. luminescens* (38), this points to an important role of signalling during insect infection. The patterns of differentially regulated genes at the two time points indicate a general transcriptional program that allows Y. enterocolitica an adaptation to insect larvae colonized at environmental temperatures.

We provide for the first time *in vivo* evidence for an involvement of the holin/endolysin cassette in Tc release and thus in nematocidal and insecticidal of *Y. enterocolitica* strain W22703. The dual lysis cassette is highly conserved in the genomes of *Yersinia* spp. where it is localised between genes encoding Tc subunits, and is also present in *P. luminescens* (9). This finding pointed to a functional role in the insecticidal activity of these bacteria (31). In *Y. pestis*, it was postulated that the release of the Tc is mediated by a type III secretion system (T3SS) (39), but this hypothesis was recently refuted (40). This is in line with the fact that a T3SS is lacking in strain W22703.

So far, there are only few examples for bacterial toxins that are released into the environment by a phage-related factors. The holin-like protein TcdE was shown to be required for *Clostridioides difficile* toxins TcdA and TcdB secretion, but TcdE seems not to affect membrane integrity of *C. difficile* as toxin release is not related to bacterial cell lysis (41, 42). A recently identified endolysin, TcdL, was demonstrated to interact with TcdB, thus providing more insight into the toxin transport mechanism (43). A further example for a correlation between phage lytic genes and toxicity is the putative coupling of the λ phage lytic cycle and the release of the phage-encoded toxin Stx from Shiga toxin producing *E. coli* (44, 45). An N-acetyl-ß-D-muramidase similar to phage endolysins was shown to be essential for *Salmonella* typhoid toxin secretion (46). A holin has not been identified to play a role in this mechanism, and the toxin components are probably secreted through the inner membrane into the periplasm via the sec-dependent route (47). In *Serratia marcescens*, a holin and endopeptidase cassette were identified to be required for the secretion of a chitinase(48). More examples were reviewed recently (49).

The holin/endolysin activity may allow the passage of the Tc through the inner membrane and the peptidoglycan layer. Given that no export signature pointing to the presence of a signal peptide for the sec-dependent pathway could be found in the toxin subunit sequences of *Y. enterocolitica*, it might be assumed that the holin acts as a pore for both the endolysin and the Tc proteins. The Tc proteins might then exert their toxic activity by being bound on the surface of the bacterial cells, as demonstrated for both YitA and YipA of *Y. pestis* (50), followed by direct contact with target cells or by a release of the enzymatically active subunit C or of the whole Tc. The frequent neighbourhood of phage-related lysis factors to bacterial toxins and other secreted factors supports the hypothesis that protein release upon the activities of a dual lysis cassette evolved multiple times and defines a more widespread mechanism that was proposed to be termed the type 10 secretion system (49). Alternatively, the Tc may be released upon cell lysis of a partial subpopulation of *Y. enterocolitica.* A detailed model of the molecular mechanism underlying the Tc release from *Y. enterocolitica*, however, remains to be elucidated.

## Conclusion

Our data demonstrate that *Y. enterocolitica* is able to colonize the midgut of *G. mellonella* and then to proliferate to high cell densities within the hemolymph of the larvae. The bacterial pathogen was shown to adapt its physiological and biochemical properties to the larvae environment within a few hours after infection. The successful infection not only depends on the insecticidal Tc, but also requires the activity of a phage-related lysis cassette.

## Materials and Methods

### Bacterial strains, plasmids and growth conditions

Bacterial strains and plasmids used in this study are listed in **Table S2**. All cultures were grown in Luria-Bertani (LB) broth (10 g/l tryptone, 5 g/l yeast extract, 5 g/l NaCl), in minimal medium consisting of M9 medium supplemented with 2 mM MgSO4, 0.1 mM CaCl2 and 55.5 mM (1% w/v) glucose or on LB agar (LB broth supplemented with 1.5% agar). *E. coli* was grown at 37°C, and *Y. enterocolitica* at 30°C or as indicated. For growth studies, overnight cultures were diluted 1:1,000 in 200 ml flasks with 50 ml medium and incubated under vigorous shaking until cells reached stationary phase, or in microtitre plates with 200 μl medium per well. If appropriate, kanamycin (50 μg ml^-1^), streptomycin (50 μg ml^-1^), chloramphenicol (20 μg ml^-1^), tetracycline (12 μg ml^-1^), nalidixic acid (20 μg ml^-1^) or arabinose as indicated were added to the media.

### General molecular techniques

DNA manipulation and isolation of chromosomal DNA was performed according to standard procedures (51), or to the manufacturer’s protocol. Polymerase chain reactions (PCR) were carried out as described recently (13). Chromosomal DNA (100 ng) was used as template for PCR amplification.

### Construction of mutants and plasmids

The construction of non-polar deletion mutants is exemplified here for the lysis cassette. Two fragments of 984 bp and 751 bp were amplified with the oligonucleotide pairs Hol.delF1/Hol.delR1 and Endo.delF2/Endo.delR2, and ligated *via* the introduced EcoRI sites. Following nested PCR with the oligonucleotides Hol.nestedAB and Endo.nestedCD and the ligation mixture as a template, the resulting fragment was cloned into pKNG101 *via BamHI*, giving rise to pKNG101ΔHE. The resulting construct was transformed from SM10 into W22703 by conjugation, and a selection procedure was performed as described recently (11). Streptomycin-sensitive clones were screened by PCR to identify mutant W22703 ΔHE. The deletion was confirmed by sequencing. To complement W22703 ΔHE, the complete coding sequence of the *holY*/*elyY* cassette and 250 nucleotides of its upstream sequence were amplified and cloned into pACYC184 *via* EcoRI in DH5α. In the resulting plasmid pACYC184-HE, the direction of *holY/elyY* transcription corresponds to that of the disrupted plasmid gene encoding the chloramphenicol acetyltransferase. Gene *tccC* was cloned into pBAD33 *via SacI* and *PstI.* All oligonucleotides used here are listed in **Table S3**.

### Bioassays

Larvae of *G. mellonella* were obtained from the Klee-Gartencenter + Zoo (Jena, Germany), and stored for less than one week at 15°C. Bacterial strains were grown to early stationary phase (optical density at 600 nm [OD_600_] ~ 1.35) at 20°C (*Yersinia* spp.), or at 37°C (DH5α). Larvae of 2-3 cm length and 120-150 mg in weight were used. Before application, the galleries were placed on ice for about 10 min as a light anesthesia. Five μl of the respective bacterial culture or its dilution in LB medium were applied orally using a Hamilton syringe (Hamilton 702 RN, 25 μl) with a very small and blunt cannula (needle gauge 33). For the application, the cannula was inserted no more than 1 mm between the labrum and labium, and the liquid was applied slowly until being completely absorbed by the insect. During infection, animals were not fed to ensure similar conditions.

Infection doses were determined by plating serial dilutions of the suspensions used for oral application. Selective agar plates with LB or *Yersinia* selective medium (Schiemann CIN medium, Oxoid, Wesel, Germany) were incubated at 30°C for 24 h. Infected larvae were incubated up to nine days in the dark at the temperature indicated, and the numbers of killed and alive larvae were enumerated each day. Larvae were considered dead if they failed to respond to touch. The TD50 was calculated using the dose-response curve (drc) package of the R software. To recover bacteria from the larvae, the larvae were surface sterilized with 70% ethanol, washed in H2O and homogenized in a mortar. The homogenous mass was suspended into 1 ml LB, rigorously shaken with a vortex, and centrifuged at 500 rpm for 30 sec. CFU were enumerated as described above. Larvae not containing cells due to imperfect application or pathogen clearance were included in the calculation.

To prepare the gut of the insects, the larvae were incubated for 10 min on ice and then placed in PBS buffer cooled to 4°C for the subsequent intestinal preparation. The larvae were carefully opened making an incision along the coronal plane from anterior to posterior end using micro scissors and Dumont forceps (Manufactures D’Outils Dumont SA, Montignez, Switzerland). The entire innards were removed and the digestive tract was dissected, removing all of the fat and organ appendages. For further analysis the prepared gut was placed in a new tube containing PBS or RNA*later* (Thermo Fisher Scientific Life Technologies GmbH, Darmstadt, Germany).

### Anatomic preparation of the digestive tract of *G. mellonella*

A solution of 2% methylene blue in distilled water was orally applied, and excessive dye was removed by washing. The main body cavity was opened from the head to tail along the ventral midline, and the cuticula was detached and pinned to a styrofoam plate. Under a binocular, the fat body and other organs were separated from the digestive tract. Oral cavity and anus were circumcised and the digestive tract removed.

### Isolation of *Y. enterocolitica* cells from *G. mellonella*

Twenty-four hours after infection, the hemolymph of *G. mellonella* was taken with a sterile insulin syringe to a volume of 100 μl, transferred into Eppendorf tubes with 500 μl resuspension buffer [1 × PBS, 0.5% biotin-free BSA, 10% (v/v) RNAlater (Thermo Fisher Scientific, Langenselbold, Germany)], and stored at 4°C for up to 24 h until further processing. After centrifugation at 9000 × g for 5 min, the pellet was resuspended and incubated in 750 μl resuspension buffer for 10 min at 4 ° C under shaking. The suspension contained a Yersinia-specific antibody (anti-Y. *enterocolitica* O:9 mouse monoclonal, FITC, PROGEN Biotechnik GmbH, Heidelberg, Germany) that was diluted 1:250. The sediment obtained by centrifugation at 9,600 × g for 2 min was washed with cold separation buffer (1 × PBS, 0.5% biotin-free BSA, 2 mM EDTA pH 7.4) containing 10% RNAlater, centrifuged, resuspended in the same buffer and incubated with 10 μl of streptavidin-coupled magnetic beads for 10 min. The washing and centrifugation steps from above were repeated, and the sediment was resuspended in 500 μl separation buffer containing 10% RNAlater. *Y. enterocolitica* cells were separated using a MACS cell separation system with a LS column (Miltenyi Biotech, Aubum, CA, USA) according to the manufacturer’s instructions.

### RNA isolation

RNA was extracted and purified from 1 ml of a *Y. enterocolitica* suspension isolated from an *in vitro* bacterial culture or from 100 μl *G. mellonella* hemolymph by immunomagnetic separation (52) using a MACS cell separation system with a LS column (Miltenyi Biotech, Aubum, CA, USA) according to the manufacturer’s instruction, and stored in RNAlater (Thermo Fisher Scientific). The bacterial pellet was homogenized with PBS, and centrifuged for 5 min at 4°C and 9000 × g. For further cell lysis, 200μl lysozme (3 mg/ml) were added and vortexed vigorously at 1,300 rpm for 15 min. After the lysate was transferred to a new tube with 50 mg of 0.1 mm Zirconia beads (Biospec, Bartlesville, U.S.A.), 700 μl of RLT buffer were added and the mixture was vortexed three times for 45 sec. After a further centrifugation step, the supernatant was transferred to a new tube, mixed with 470 μl of 100% ethanol, applied to the column, and processed further in the RNeasy^®^ Mini Kit, according to the manufacturer’s instructions (QIAGEN GmbH, Hilden, Germany). RNA quality was assessed using a 2100 Bioanalyser (Agilent, Waldbronn, Germany).

### Transcriptome analysis

Whole-transcriptome RNA library preparation with isolated RNA was performed as described (53). Briefly, ribosomal RNAs were depleted using the Ribominus Transcriptome isolation Kit (Invitrogen, Darmstadt, Germany), and RNA was fragmented *via* a Covaris sonicator. Following de- and rephosphorylation, the TruSeq Small RNA Sample Kit (Illumina, Munich, Germany) was used for RT-PCR and gel purification of the resulting cDNAs. Libraries were then diluted and sequenced on a MiSeq sequencer (Illumina, Munich, Germany) using a MiSeq Reagent Kit v2 (50 cycles), resulting in 50 bp single-end reads. Illumina FASTQ files were mapped to the reference genome of *Y. enterocolitica* W22703 (Accession number PRJEA59689; (10)) using Bowtie for Illumina implemented in Galaxy (54, 55). Artemis was used to visualize and calculate the number of reads mapping on each gene (56, 57). Gene counts of each library were normalized to the smallest library in the comparison, and RPKM (reads per kilobase per million mapped reads) values were calculated. Fold changes between the different conditions were determined.

### Histology

*G. mellonella* larvae were immobilized at 4°C, pinned on cork discs in a stretched position and placed in 4% neutral buffered formaldehyde at least twelve hours for fixation. Larvae were removed from the cork discs and placed in 3 cm × 2,5 cm × 0,5 cm metal molds filled with 10% liquid agarose. They were oriented by forceps in a straight position with the legs up, and the agarose was allowed to solidify for 30 min at 4°C. After removing excessive agarose, larvae were dissected with a razor blade along the body midline in equal halves using mouth, tail and legs for orientation. Both parts were placed with the cut surface downwards in cassettes. For paraffin embedding, formalin was removed by rinsing in tap water, tissues were dehydrated in graded ethanols followed by xylene, infiltrated with paraffin type 1 (Richard Allan Sci., Michigan, USA) in an automatic vacuum infiltration processor (Tissue-Tek VIP^®^ 6, Sakura, Staufen, Germany) and embedded in paraffin type 6 (Richard Allan Sci., Michigan, USA). Consecutive, 2 μm paraffin sections made by an Epredia™ HM 355S automatic microtome (Fisher Scientific GmbH, Schwerte, Germany) were collected on charged glass slides and dried at 37°C for 24 to 48 h. Deparaffinised tissue section were stained with hemalaun and eosin for morphological orientation and identification of tissue lesions.

### Immunofluorescence

To detect *Y. enterocolitica* by fluorescent-labelled antibodies, consecutive tissue sections were deparaffinized in xylene and graded ethanols, and antigen-demasked by digestion with 0.1% trypsin for 20 min at 37°C. After washing in PBS, a fluorescent-labelled anti-Y. *enterocolitica* O:9 mouse monoclonal antibody (FITC; PROGEN Biotechnik GmbH, Heidelberg, Germany) was diluted 1:250 in PBS containing 3% BSA and applied for 1 h at R. T. An anti-E. *coli* polyclonal antibody (FITC) diluted 1:100 in PBS was used in controls (Genetex, Biozol Diagnostica Vertrieb GmbH, Leipziger Straße 4, 85386 Eching, Germany). After a further washing step, tissue sections were covered-slipped with ProLong™ Diamond Antifade Mountant (Thermo Fisher Scientific, Darmstadt, Germany). Microscopic slides were evaluated on a Zeiss Axio Imager 2 (Carl Zeiss Microscopy GmbH, Göttingen, Germany) using a dual emission filter set, allowing the simultaneous detection of FITC-marked bacteria in the green channel and imaging of the insect tissue through their own fluorescence in the red channel.

### Immunohistochemistry

Analogous to immunofluorescence, the dewaxed tissue sections were digested with trypsin (0.05 g trypsin, 0.66 g CaCl_2_ × 2H2O, 50 ml dH_2_O, pH 7.8) for 20 min at 37°C in a water bath. After washing for 5 min in PBS, the primary antibody (anti-*Y. enterocolitica* O:9 mouse monoclonal, FITC, PROGEN Biotechnik GmbH) was diluted 1:250 in PBS containing 3% BSA and incubated for 1 h at RT. After three washing steps of 5 min each in PBS, the secondary antibody (HRP-Polymer-anti-mouse, Zytomed Systems GmbH, Berlin, Germany) was diluted 1:2 in PBS containing 3% BSA and incubated for 1 h at RT. After incubation for 5-10 min with 3,3’-diaminobenzidine, counterstaining with hematoxylin took place. Finally, the stained tissue sections were covered with Kaiser’s glycerine gelatin and analysed via the Zeiss Axio Imager 2.

### Giemsa staining of hemocytes

*G. mellonella* larvae were picked up, pricked on the aorta in the abdominal area with a sharp cannula and the escaping hemolymph fluid was used for a smear on a microscopic slide. The preparations were dried for about 10 min at RT, followed by fixation for 10 min with methanol or acetone. The sections were placed in dH2O twice for 2 min and stained in the Giemsa solution (5 ml Giemsa stock solution in 80 ml dH_2_O) for 30 min. After staining, the preparation was differentiated with dH_2_0 (and a few drops of acetic acid) until it was mauve. Another differentiation followed in 96% ethanol until no more clouds of color came off. In the last steps, the slides were washed 3 × 2 min in isopropanol, placed 2 × 2 min in xylene and covered with Eukitt^®^ Quick-hardening mounting medium (Merck KGaA, Darmstadt, Germany). The microscopic slides were evaluated on a Zeiss Axio Imager 2 (Carl Zeiss Microscopy GmbH).

## Acknowledgments

We thank the Deutsche Forschungsgemeinschaft (grant FU 375/4-1, 2 to TMF) for financial support of this study, Jana Jagiela und Lisa Wolf for technical assistance, and Marcus Pfau for photographic work.

## Supplementary Material

**Figure S1.** The HE lysis cassette within Tc-PAI_*Ye*_.

**Table S1.** Heat map of differentially regulated *Y. enterocolitica* genes *in vivo.*

**Table S2.** Bacterial strains and plasmids used in this study.

**Table S3.** Primers.

