## Supplementary material for "Dissecting the invasion of *Galleria mellonella* by *Yersinia enterocolitica* reveals metabolic adaptations and a role of a phage lysis cassette in insect killing": Table S2. Strains and plasmids.docx

**Table S2.** Strains and plasmids used in this study.

| **Strains** | | **genotype/relevant features** | **reference or source** |
| --- | --- | --- | --- |
| *E. coli* | |  | |
|  | DH5α | F^-^ φ80Δ*lac*ZΔM15 Δ(*lac*ZYA-*arg*F) U169 *rec*A1 *end*A1 *hsd*R17(r_k_^-^, m_k_^+^) *pho*A *sup*E44 λ- *thi*-1 *gyr*A96 *rel*A1 | Invitrogen, Karlsruhe, Germany |
| *Y. enterocolitica* | |  | |
|  | W22703 | Nal^R^, Res^-^ Mod^+^, pYV^-^ | {Cornelis, 1975 #382} |
|  | W22703 Δ*tcaA* | W22703 mutant with a non-polar deletion of *tcaA* | {Bresolin, 2006 #558} |
|  | W22703 ΔHE | W22703 mutant with a non-polar deletion of *holY and elyY* | This study |
|  | W22703 Δ*tccC* | W22703 mutant with a non-polar deletion of *tccC* | This study |
|  | W22703 Δ*tcaR2* | W22703 mutant with a non-polar deletion of *tcaR2* | {Starke, 2014 #779} |
|  | W22703 *tcaA*::*rfp* | Gene *rfp* fused immediately behind *tcaA* via chromosomal insertion of pUTs-`*tcaA*::*rfp* | {Starke, 2014 #779} |
|  | W22703 P_HE_::*rfp* | Gene *rfp* fused to the HE promoter via chromosomal insertion of pUTs-P_HE_::*rfp* | This study |
| **Plasmids** | |  | |
| pKNG101 | | Conditionally replicating vector; R6K origin, mobRK2 transfer origin, sucrose-inducible *sacB*, Str^R^ | {Kaniga, 1991 #555} |
| pUTs-*lux*(Cm) | | Cm^R^, transposase-negative derivative of pUT mini-Tn*5 luxCDABE* Km2; suicide plasmid in *pir* negative strains | {Starke, 2013 #763} |
| pKRG9 | | Derivative of suicide vector pGP704 {Miller, 1988 #236}; *ori* R6K, mob^+^ (RP4), Cm^R^, Amp^S^ | Creatogen, Augsburg, Germany |
| pKD4 | | *pir* dependent, FRT sites, Kan^R^ | {Datsenko, 2000 #711} |
| pKD119 | | Lambda red helper plasmid, Tet^R^ | {Datsenko, 2000 #711} |
| pCP20 | | FLP recombinase plasmid, Cm^R^, Amp^R^ | {Datsenko, 2000 #711} |
| pUTs-*rfp*(Cm) | | As above, *luxCDABE* exchanged with *rfp* | {Starke, 2013 #763} |
| pUTs-`*tcaA*::*rfp* | | Last 500 bp of *tcaA* cloned in front of *rfp* within plasmid pUTs-*rfp*(Cm) | {Starke, 2014 #779} |
| pUTs-P_HE_::*rfp* | | 500 bp upstream of *hlyY* cloned via *Sac*I and *Kpn*I in front of *rfp* within plasmid pUTs-*rfp*(Cm) | This study |
| pACYC184 | | p15A origin, Cam^R^, Tet^R^, | {Chang, 1978 #556} |
| pACYC-*tcaA* | | pACYC184 with an *Eco*RI fragmentcontaining *tcaA* and its promoter region, Cam^S^) | {Bresolin, 2006 #558} |
| pACYC-HE | | Gene *holY* and *elyY* including a 500 bp upstream region cloned via *Eco*RI into pACYC184, Cam^S^ | This study |
| pBAD33 | | Expression vector with arabinose-inducible promoter, Cam^R^ | {Guzman, 1995 #732} |
| pBAD33-*tccC* | | Gene *tccC* cloned into pBAD33 *via* *Sac*I and *Pst*I | This study |
