## Supplementary material for "Dissecting the invasion of *Galleria mellonella* by *Yersinia enterocolitica* reveals metabolic adaptations and a role of a phage lysis cassette in insect killing": Table S3. Primer.docx

Supplementary Table S2: Oligonucleotides used in this study.

| name | sequence 5´-3´ | purpose |
| --- | --- | --- |
| Hol.delF1 | cagaacttcaggatggc | W22703 ΔHE |
| Hol.delR1 | gagaattcgaggattaagaagtccattt | W22703 ΔHE |
| Endo.delF2 | ggcaacttaactcacga | W22703 ΔHE |
| Endo.delR2 | gagaattcccctattttgaattaccca | W22703 ΔHE |
| Hol.nestedAB | cgggatccgcactgaggtacaacg | W22703 ΔHE |
| Endo.nestedCD | cgggatccgcgttatagcggttgc | W22703 ΔHE |
| HEkomplF | ccggaattcgtgatgacagcgctcg | pACYC-HE |
| HEkomplR | ccggaattctttagacatggtaattttcc | pACYC-HE |
| TccC_delR1 | ccggaattcgtgggagtttgttcacaaag | W22703 Δ*tccC* |
| TccC_delF1 | atgacagccctcagtttc | W22703 Δ*tccC* |
| TccC_delF2 | ccggaattcgcagtagaacgtagtgaag | W22703 Δ*tccC* |
| TccC_delR2 | aatacgccgtgtaaaccc | W22703 Δ*tccC* |
| TccC.nestedF | cgggatcccgcgctctgactctaag | W22703 Δ*tccC* |
| TccC.nestedR | cgggatccagggcagattgaatggtg | W22703 Δ*tccC* |
| TccC_SacI_F2 | cgatgagctcatgtctaaaacgtcatttg | pBAD33-*tccC* |
| TccC_PstI_R2 | aactgcagctaatttaataagtgcggg | pBAD33-*tccC* |
| PholF | cgatgagctccctcatgccatgaccg | pUTs-P_HE_::*rfp* |
| PholR | cggggtacctttgtttacctatcattgttg | pUTs-P_HE_::*rfp* |
