## Supplementary figures and images for "Dissecting the invasion of *Galleria mellonella* by *Yersinia enterocolitica* reveals metabolic adaptations and a role of a phage lysis cassette in insect killing"

### Figure S1 The HE lysis cassette within Tc-PAIYe.pptx

#### Slide 1
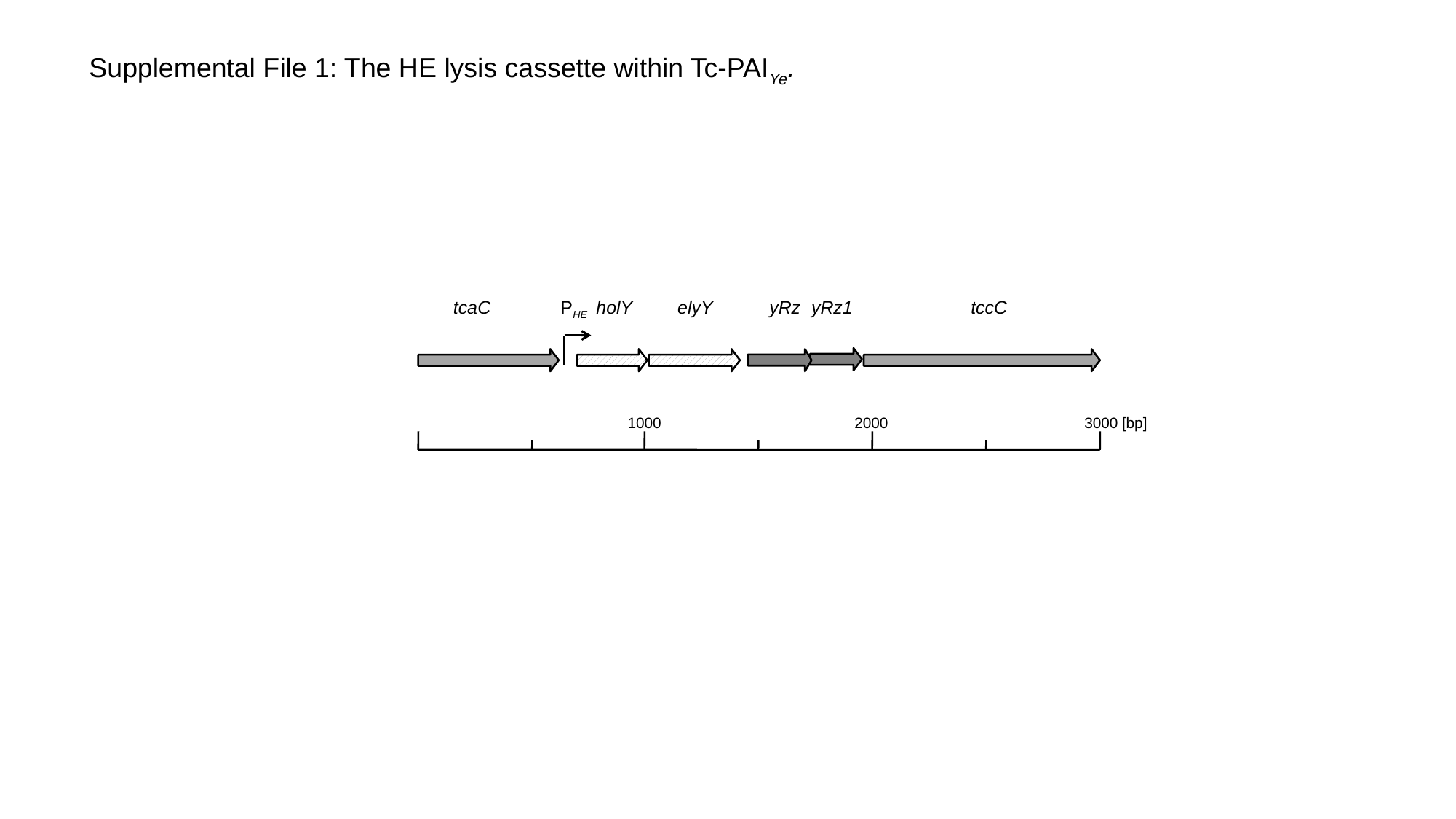

Supplemental File 1: The HE lysis cassette within Tc-PAI_Ye_.
tcaC
P_HE_
holY
elyY
yRz
yRz1
tccC
1000
2000
3000
[bp]
